# Portrait of a genus: genome sequencing reveals evidence of adaptive variation in *Zea*

**DOI:** 10.1101/2022.06.03.494450

**Authors:** Lu Chen, Jingyun Luo, Minliang Jin, Ning Yang, Xiangguo Liu, Yong Peng, Wenqiang Li, Alyssa Philips, Brenda Cameron, Julio Bernal, Rubén Rellán-Álvarez, Ruairidh JH Sawers, Liu Qing, Yuejia Yin, Xinnan Ye, Jiali Yan, Qinghua Zhang, Xiaoting Zhang, Shenshen Wu, Songtao Gui, Wenjie Wei, Yuebin Wang, Yun Luo, Chengling Jiang, Min Deng, Min Jin, Liumei Jian, Yanhui Yu, Maolin Zhang, Xiaohong Yang, Matthew B. Hufford, Alisdair R. Fernie, Marilyn L. Warburton, Jeffrey Ross-Ibarra, Jianbing Yan

## Abstract

Maize is a globally valuable commodity and one of the most extensively studied genetic model organisms. However, we know surprisingly little about the extent and potential utility of the genetic variation found in the wild relatives of maize. Here, we characterize a high-density genomic variation map from 744 genomes encompassing maize and all wild taxa of the genus *Zea*, identifying over 70 million single nucleotide polymorphisms (SNPs) and nearly 9 million Insertion/Deletion (InDel) polymorphisms. The variation map reveals evidence of selection within taxa displaying novel adaptations to traits such as waterlogging, perenniality and regrowth. We focus in detail on adaptive alleles in highland teosinte and temperate maize and highlight the key role of flowering time related pathways in highland and high latitude adaptation. To show how this data can identify useful genetic variants, we generated and characterized novel mutant alleles for two flowering time candidate genes. This work provides the most extensive sampling to date of the genetic diversity of the genus *Zea*, resolving questions on evolution and identifying adaptive variants for direct use in modern breeding.

## Introduction

Global crop production is currently insufficient to meet the anticipated demands of a growing human population^1,2^. Climate change is affecting crop production in many areas, further exacerbating this problem^3^, and projected shifts in temperature and precipitation will lead to further declines in productivity for many major crops^4^. New varieties displaying both higher yield and the better adaptation to diverse environments are thus urgently needed to increase crop productivity under changing climate scenarios^5,6^.

Maize (*Zea mays* subsp. *mays*) is one of the world’s most widely grown crops, with an annual global production of over 1.1 billion tons in 2018 (FAOSTAT, 2020). Native American peoples domesticated from the wild grass *Zea mays* subsp. *parviglumis* (hereafter *parviglumis*) approximately 9,000 years ago in the southwest of Mexico^7,8^. Population genetic analyses largely agree that maize underwent a substantial population bottleneck during domestication^9–12^, reducing the genetic diversity available for adaptation. Although maize rapidly spread from its center of domestication across a wide range of environments, successful adaptation required hundreds or thousands of years^13^. As global populations increase and climate change accelerates, unprecedented maize yield losses are projected to become commonplace in most maize-producing regions^5,14,15^. To facilitate adaptation to these new challenges, breeders will need to maximize use of the genetic diversity at their disposal, looking beyond modern elite lines to traditional cultivated varieties and locally adapted wild relatives^16^.

The wild congeners of maize, collectively called teosintes, are annual and perennial grasses native to Mexico and Central America (Fig. 1a). They are adapted to a diverse range of environments, from hot, humid, subtropical regions of Central America to cold, dry, high elevations of the Mexican Central Plateau^17,18^. Teosintes exhibit biotic and abiotic adaptations absent in modern maize and humid high elevations in central Western Mexico and the Huehuetenango region of Guatemala^17–19^, providing a wealth of genetic diversity that could be utilized in modern breeding. A recent example is a large-effect allele for leaf angle identified in teosinte that was lost during maize domestication^20^. CRISPR-Cas9 editing of maize to mimic the teosinte allele resulted in a ~20% yield increase in modern hybrids grown at high density. Other studies have used genetic mapping to capitalize on teosinte alleles for nutrition^21,22^, adaptation to extreme environments^23,24^, and disease resistance^25–27^. Population genetic evidence suggests that diverse alleles from the teosinte *Zea mays* subsp. *mexicana* (hereafter *mexicana*) played an important role in allowing maize to adapt to arid highland conditions^28–30^.

**Fig. 1.**
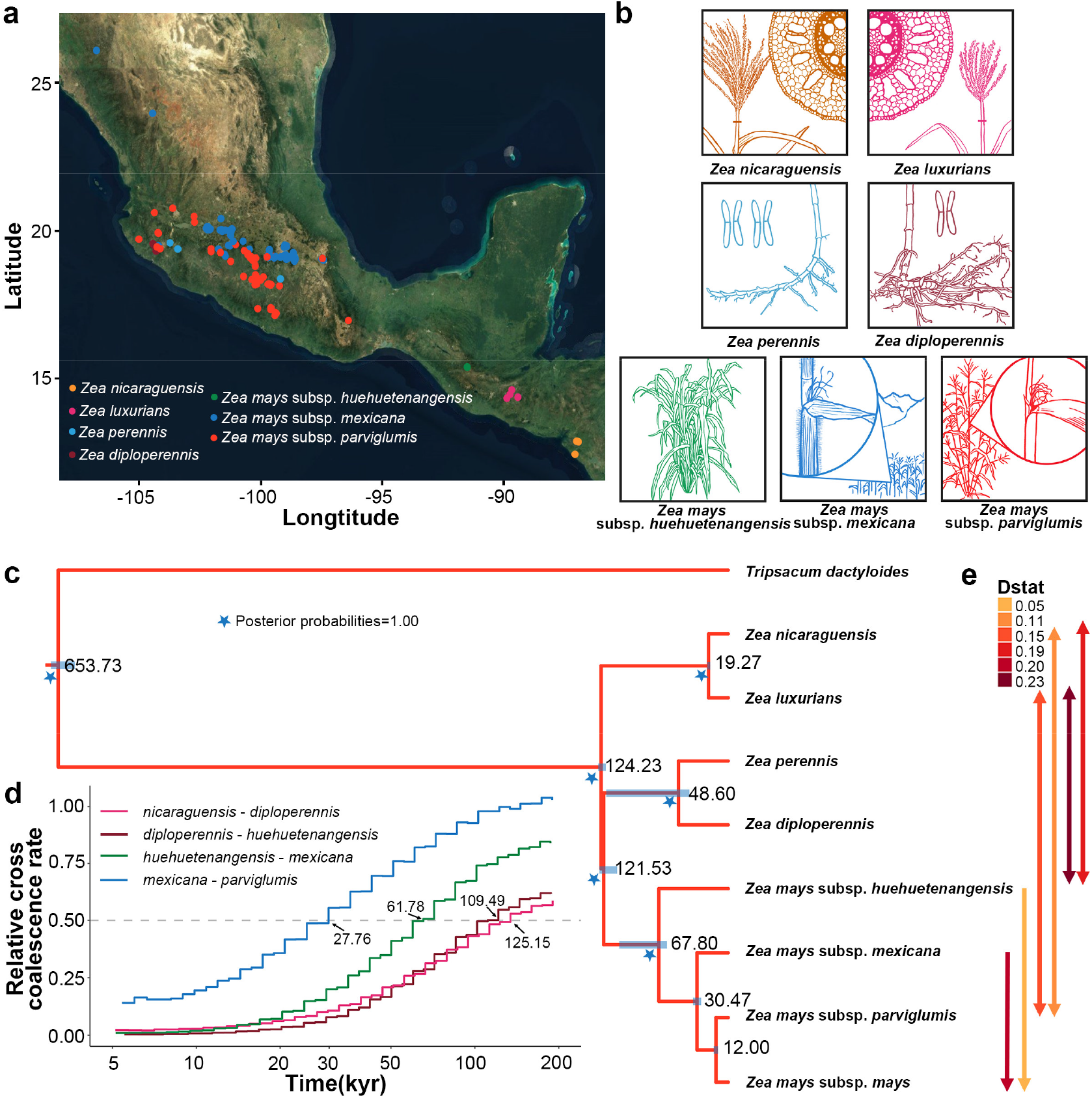
Phylogeny of *Zea* genus. **a**, Geographical distribution of collected teosintes, taxa were identified and colored based on morphology. **b**, Morphological characteristics of teosinte (Credit to Dr. Andi Kur). *nicaraguensis* and *luxurians* are distinguished from other teosinte based on aerenchyma in their stems which aerate roots during submergence, while *nicaraguensis* has a more robust tassel than *luxurians. perennis* is a recent autotetraploid of *diploperennis*; the rhizomatous root systems of these perennial taxa distinguish them from other teosintes. The Mexican annual teosintes *parviglumis* and *mexicana* are distinguished from each other based on the presence of macro-hairs and pigment along their stems, two traits that are linked to highland adaptation. **c**, Divergence times (in thousands of years before present) estimated from the multispecies coalescent (MSC) model. Blue bars indicate the 95% highest posterior density (HPD) intervals. The star indicates nodes with posterior probability of 1. Edge widths reflect estimates of effective population size (Supplementary Table 4). **d**, Rates of cross-population coalescence among teosinte species. Curves were computed using four phased haplotypes. **e**, Introgression among taxa. Arrows indicate the taxa involved (one-way arrow indicate unidirectional introgression, two-way arrow indicate bidirectional introgression), and arrow color shows the value of Patterson’s D-statistic (Supplementary Table 5).

Despite the potential for teosinte to contribute to breeding and adaptation of cultivated maize, we know relatively little about the genetic diversity and history of these taxa. Estimates of the age of the genus vary substantially^31–35^, and the phylogenetic relationship of several taxa is debated or unknown^17,36–38^. Considerable cytological diversity is found within the genus, and transposable element variation^39–41^ and large inversions^42–47^ have been documented as well. Moreover, common garden studies have demonstrated that phenotypic differentiation in both teosinte and maize landraces is the result of local adaptation^48,49^. Low density genotyping or pooled sequencing approaches in *parviglumis* and *mexicana* have identified a number of candidate loci related to soil, climate, and disease resistance, highlighting the importance of inversions^46,50,51^. However, for most taxa in *Zea*, their potential as sources of useful diversity in maize remains poorly understood.

Here, we present a genus-wide resource of genome-scale genetic diversity in *Zea*. We resequenced 237 teosinte accessions, including all seven taxa of teosinte, and combined these data with sequences from 507 maize inbred lines. Our analyses reveal a detailed phylogeny and demography of the genus *Zea*, identify substantial novel genetic diversity, and expand our understanding of adaptation in the genus *Zea*. We predict these resources will substantially facilitate the efficient use of diverse *Zea* taxa in modern maize breeding and improvement.

## Results

### The diversity map and phylogeny of the genus *Zea*

We resequenced 237 teosinte accessions encompassing all described species and subspecies in the genus *Zea* (Fig. 1a, b) to an average depth of 22x, and combined these data with genome resequencing data from 507 cultivated maize inbred lines representing both temperate and tropical regions^52^ (Supplementary Table 1). To ensure the quality of this new *Zea* diversity map, we used a set of strict filtering conditions (Methods) and identified a final set more than 70M SNPs and nearly 9M insertion/deletions (InDels) (Supplementary Table 2), with nearly 80% of SNPs segregating as rare variants (MAF<0.05) (Supplementary Fig. 1). Both classes of variants appeared enriched in genic and regulatory regions (30% of SNPs and 45% of InDels in 14% of the genome), likely reflecting difficulties in read mapping in repetitive regions of the genome. We validated a subset of genic SNPs using Sanger sequencing, with median concordance between datasets >95% and reasonable false positive and false negative rates (both ~5% on average) for non-reference alleles (Supplementary Table 3). Based on population structure analysis, samples with greater than 60% ancestry in a single group were clustered into *parviglumis* (*n*=70), *mexicana* (*n*=81), *Zea mays* subsp. *huehuetenangensis* (*n*=5; hereafter, *huehuetenangensis*), *Zea diploperennis* (*n*=20; hereafter, *diploperennis*), *Zea perennis* (*n*=19; hereafter, *perennis*), *Zea luxurians* (*n*=14; hereafter, *luxurians*), *Zea nicaraguensis* (*n*=14; hereafter, *nicaraguensis*), 210 tropical maize and 280 temperate maize (Supplementary Fig. 2a,b and Supplementary Table 1). Principal component analysis of these lines was in strong concordance with population structure results (Supplementary Fig. 2c).

We inferred phylogenetic relationships for the genus *Zea* under the multispecies coalescent model^53^ (Fig. 1c); maximum likelihood phylogenies^54^ produced largely congruent results (Supplementary Fig. 3, 4). Notably, we estimated a very recent origin for the genus, splitting from its sister genus *Tripsacum* only ~650,000 years ago. This young age is especially striking given the pronounced differences in chromosome structure and sub-genome organization resulting from the two genera’s shared polyploidy event >10M years ago^55^. Within the genus, our results suggest that *nicaraguensis* likely represents a subspecies of *luxurians*, with divergence times similar to those among subspecies of *Zea mays*. The phylogeny supports earlier analysis^34^ suggesting that divergence among *Zea mays, luxurians*, and *diploperennis* was nearly contemporaneous, occurring ~120,000 years ago (95% highest posterior density (HPD) interval for *luxurians* divergence from other taxa: 125,967-127,200; Fig. 1c and Supplementary Table 4). We further estimate that *perennis* split from its diploid progenitor *diploperennis* only ~48,000 years ago (95% HPD: 38,033-119,100). Tree topologies and divergence times also support earlier analyses^56^ showing that *huehuetenangensis* is a subspecies of *Zea mays*, diverging from other annual subspecies ~68,000 years ago (95% HPD: 60,133-106,467), followed by the divergence of highland *mexicana* and lowland *parviglumis* ~30,000 years ago (95% HPD: 26,733-34,500). Our phylogeny estimates the divergence of maize from *parviglumis* at ~12,000 years, only slightly older than the earliest archaeological evidence^8^ and likely due to population structure within *parviglumis*^37,46^. Independent estimates of divergence times taken from rates of cross-coalescence^57^ between taxa are strikingly consistent (Fig. 1d).

Population genetic analysis of diversity further reveals changes in demography among taxa in *Zea*. Coalescent estimates of effective population size (*N_e_*) over time reveal the well-established bottleneck associated with maize domestication but also a continued decline in population size for the annual subsepecies *parviglumis* and *mexicana* since their divergence (Supplementary Fig. 5). All other taxa in the genus show parallel trends, with steady declines in population size until about 10,000 years ago, with more recent increases for *luxurians* and *diploperennis*. Patterns of shared derived alleles and sequence divergence both suggest a history of introgression among taxa (Fig. 1e, Supplementary Fig. 6 and Supplementary Table 5), including bidirectional admixture between *parviglumis/huehuetenangensis* and *nicaraguensis/luxurians*, and unidirectional introgression from *huehuetenangensis/mexicana* into domesticated maize, highlighting the important role of gene flow in crop adaptation^58^.

### Novel diversity in *Zea*

SNP data highlight the impressive genetic diversity present in teosinte. Despite the potential downward bias due to strict filtering parameters and read mapping to a maize reference, heterozygosity and nucleotide diversity are both higher in teosinte taxa than the much larger panel of maize lines, even among teosinte with limited geographic ranges (Supplementary Table 2 and Supplementary Fig. 7). Nearly a quarter (24%) of the SNPs and 20% of the InDels identified across all taxa are taxon-specific (Supplementary Table 2), and there are significantly more SNPs specific to each teosinte accession than maize (Fig. 2a), this tendency remains the same after choosing comparable samples in each taxon (Supplementary Fig. 8). In teosintes, a substantial proportion of taxon-specific SNPs and InDels are located in genic and regulatory regions (promoter and cis-regulatory elements^59^; Supplementary Fig. 9), suggesting the presence of biologically functional alleles with potential for improving modern maize. Differentiation (*F*_ST_) between teosinte taxa is often lower than that found between inbred maize and teosinte (Supplementary Fig. 10a), consistent with the historical reduction of diversity that occurred during modern maize breeding^60^. The annual subspecies of *Zea mays* show much faster decay of linkage disequilibrium than our diverse panel of maize inbreds (10-50Kb compared to ~200Kb; Supplementary Fig. 10b), but historical recombination in other teosintes appears to be even more limited (>500Kb).

**Fig. 2.**
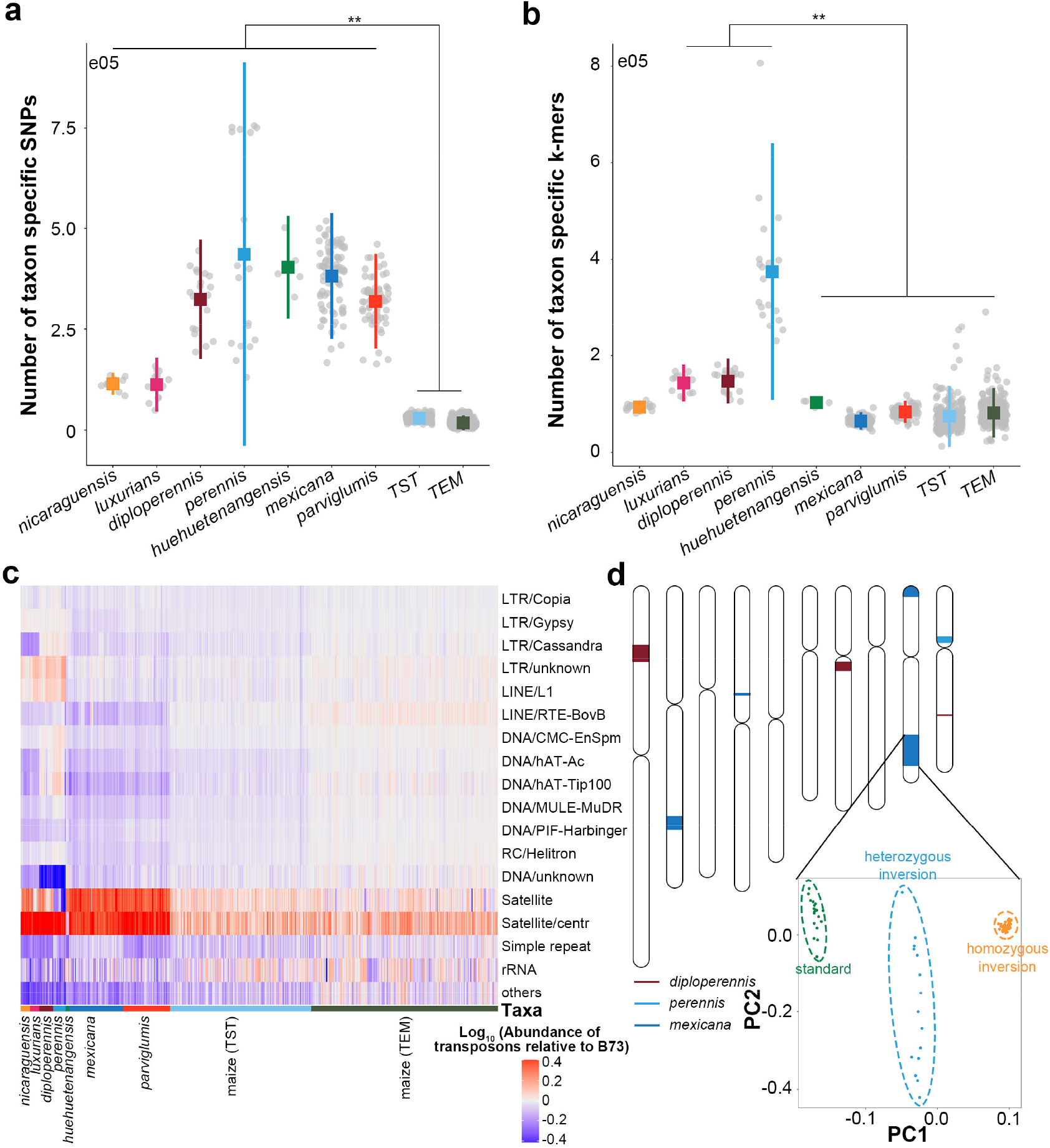
Variation in the *Zea* genus. **a**, Taxon specific SNPs and **b**, k-mers (31bp) in *Zea* genus. TST indicates tropical maize, and TEM indicates temperate maize. The significantly lines compare all teosinte to TST and TEM (**a**) or *luxurians/diploperennis/perennis* to *Zea mays* (**b**). **c**, Abundance of transposon elements relative to B73. Each column represents a sample. **d**, Distribution of inversions across the chromosomes. Each colored segment represents an inversion, with colors referring to the population in which the inversion is most prevalent (deep red: *diploperennis*; blue: *perennis*; deep blue: *mexicana*). Inset shows PCA of SNPs data from within *Inv9e*, clearly separating the three genotype classes (left: standard; middle: heterozygous inversion; right: homozygous inversion).

Short-read mapping approaches pose challenges in characterizing genetic diversity, including difficulty with repetitive sequences and reference bias. In order to circumvent some of these obstacles, we used a reference-free k-mer approach to characterize each taxon (Methods). Consistent with the reference mapping bias (~8% unmapped reads in average), most taxa showed a substantial proportion of unique k-mers (Supplementary Fig. 11a, b Supplementary Table 2). Since non *Zea mays* species have diverged from *Zea mays* more than ~120,000 years (Fig. 1c), the higher number of unique k-mers were exhibited in their genomes as expected (Fig. 2b and Supplementary Fig. 11c, d). These results not only highlight the novel genetic diversity present in teosinte but also likely point to the ongoing importance of evolutionary processes in generating and filtering diversity in traditional maize populations in Mexico^61^.

We next investigated the diversity and abundance of transposons and inversion polymorphisms in *Zea*. Transposable elements (TEs) are an important driver of shaping the structure and evolution of the genome^62^, and over 85% of the maize genome is repetitive sequence^63^. Clustering repeats from our short-read data accounted for ~74% of sequence across the genus (Supplementary Table 6), with the vast majority (60-70%) coming from LTR retrotransposon. Mapping reads from individual genomes to these clusters revealed broadly similar patterns across species, consistent with previous comparisons of *Zea mays* and *luxurians*^64^. Nonethless, we do identify a notable decrease in the percentage of Ty3 retrotransposons in *Zea mays* compared to other species, and an increase abundance of DNA transposons in *diploperennis* and *perennis* (Fig. 2c and Supplementary Fig 12).

Inversions are known to play important roles in adaptation and speciation^65,66^, and previous work has highlighted the evolutionary relevance of several large inversions in *Zea*^23,45,46,67^, including *Inv9e* in *mexicana* adaptation^46,50,51^. Multidimensional scaling of SNP diversity across the genome^68^ allowed us to identify eight large genomic regions (> 1 Mb) indicative of inversion polymorphism (Supplementary Fig. 13, Supplementary Table 7). Six of these are newly identified in the present study, and show clustering patterns delineating the three genotypes (standard; heterozygous inversion and homozygous inversion; Fig. 2d, Supplementary Fig. 14 and Supplementary Table 8).

Given previous evidence suggesting the association between inversions and soil characteristics^46^, we performed genome-wide association with nine representative soil traits (Methods) from a rich database of more than 200 soil properties^69^ (Supplementary Fig. 15a and Supplementary Table 9). *Inv9e* was significantly associated with gypsum content (0.829-1.383m) which is a representative of 29 soil properties (Supplementary Fig. 15b and Supplementary Table 9). We merged nearby significant SNPs located in *Inv9e* into two QTLs (chr9:127,017,047-127,356,295 and chr9:138,354,955-139,846,464; Supplementary Fig. 16, Supplementary Table 10). These contain 15 genes that have been functionally validated in rice or *Arabidopsis* (Supplementary Table 11) including two (*Zm00001d047667* and *Zm00001d047694*) with orthologs that have been confirmed to affect root development in rice^70,71^ and may provide clues to further explore the function of *Inv9e* in adaptation. Given that many inversions found segregating at appreciable frequency are likely adaptive in some environments^72,73^, these data argue that improved assembly and characterization of structural variants in teosinte would be a promising avenue for discovery of new functional genetic diversity.

### Signals of selection from allele frequency data

Their genetic, ecological, and life history diversity make teosintes an ideal model system for studying adaptation^17^. To identify potential targets of selection, we calculated *F*_ST_ between each teosinte taxa and cultivated maize in 5-kb sliding windows (Methods). Here, we found a high proportion of outlier windows shared between the closely related taxa (56% overlapped between *nicaraguensis* and *luxurians*; 54% overlapped between *diploperennis* and *perennis*; Supplementary Table 12, Supplementary Fig. 17). Shared genes (5,706; Supplementary Table 13) in *nicaraguensis* and *luxurians* comparisons were enriched in core cell component and reproductive system developmental processes (GO:0061458; *P*-value = 1.15E-04; FDR = 6.87E-03; Supplementary Table 14). Candidate adaptive genes (4,659; Supplementary Table 15) in *diploperennis* and *perennis* comparisons were enriched in some basic biological process and core cellular components such as nucleus (GO:0005634; *P*-value = 1.25E-12; FDR = 2.89E −10) (Supplementary Fig. 18, Supplementary Table 16).

We also identify a number of genes related with known pathways involved in meiosis^74^, QTLs in regrowth^75^ and waterlogging^76–78^ (Supplementary Table 17). These include *Zm00001d002945*, an ortholog of the *Arabidopsis* gene *AtNAC082* involved in the regulation of leaf senescence^79^, which shows high *F*_ST_ in *diploperennis* - maize and *perennis* - maize comparisons and is located in a QTL region controlling regrowth^75^. In *nicaraguensis* - maize, *luxurians* - maize comparisons, we find genes potentially involved in the response to waterlogging not only by regulating the content of ethylene and wax, but also the photosynthetic efficiency to adapt to the wetter climate in Guatemala^17^. These include *Zm00001d015637*, the maize ortholog of *AtOSP1* in *Arabidopsis*, a GDSL lipase that is required for wax biosynthesis and stomatal formation^80^. These genes highlight the value of our diversity data in identifying candidate loci of potential adaptive relevance for maize, and present a catalog of genes worth further exploration.

In addition to identifying differences among species, our extensive sampling of *parviglumis* (*n*=70), *mexicana* (*n*=81), and both temperate (*n*=280) and tropical maize (*n*=210) accessions allowed investigation of more recent adaptation to highlands and high latitudes. Both the high elevation and high latitude reflects a climate of lower temperate and longer light period, and previous work identified evidence of convergent selection between temperate maize and its broadly-distributed temperate relative *Tripsacum*^81^. Here, we extended this comparison to investigate convergence between temperate maize and high elevation adapted teosinte (*mexicana*). We applied a composite likelihood genome-scan (see Methods) for selection between *mexicana* vs *parviglumis* and temperate vs tropical maize (Fig. 3a, b and Supplementary Table 18, 19). We found significant overlap in selected windows (*P* = 0.047; 14.7% higher than permutations; Supplementary Fig. 19a), but less overlap than expected in candidate genes (*P* = 0.97; 27% less than permutation). Notably, however, ~90% of selected windows in both comparisons were found in noncoding regions of the genome, suggesting adaptation may predominantly have targeted regulatory regions. To test for convergence in regulatory adaptation, used RNA-seq from the shoot base of *parviglumis*, *mexicana* and tropical and temperate maize to search for changes in gene expression. We identified 595 genes differentially expressed between *mexicana* and *parviglumis* (Supplementary Table 20) and 437 genes differentially expressed between tropical and temperate maize (Supplementary Table 21), with significant overlap between the two lists (*P* = 0.006; 102% higher than permutations; Supplementary Fig. 19b). Those results may point to the importance of convergent regulatory evolution in maize and teosinte local adaptation.

**Fig. 3.**
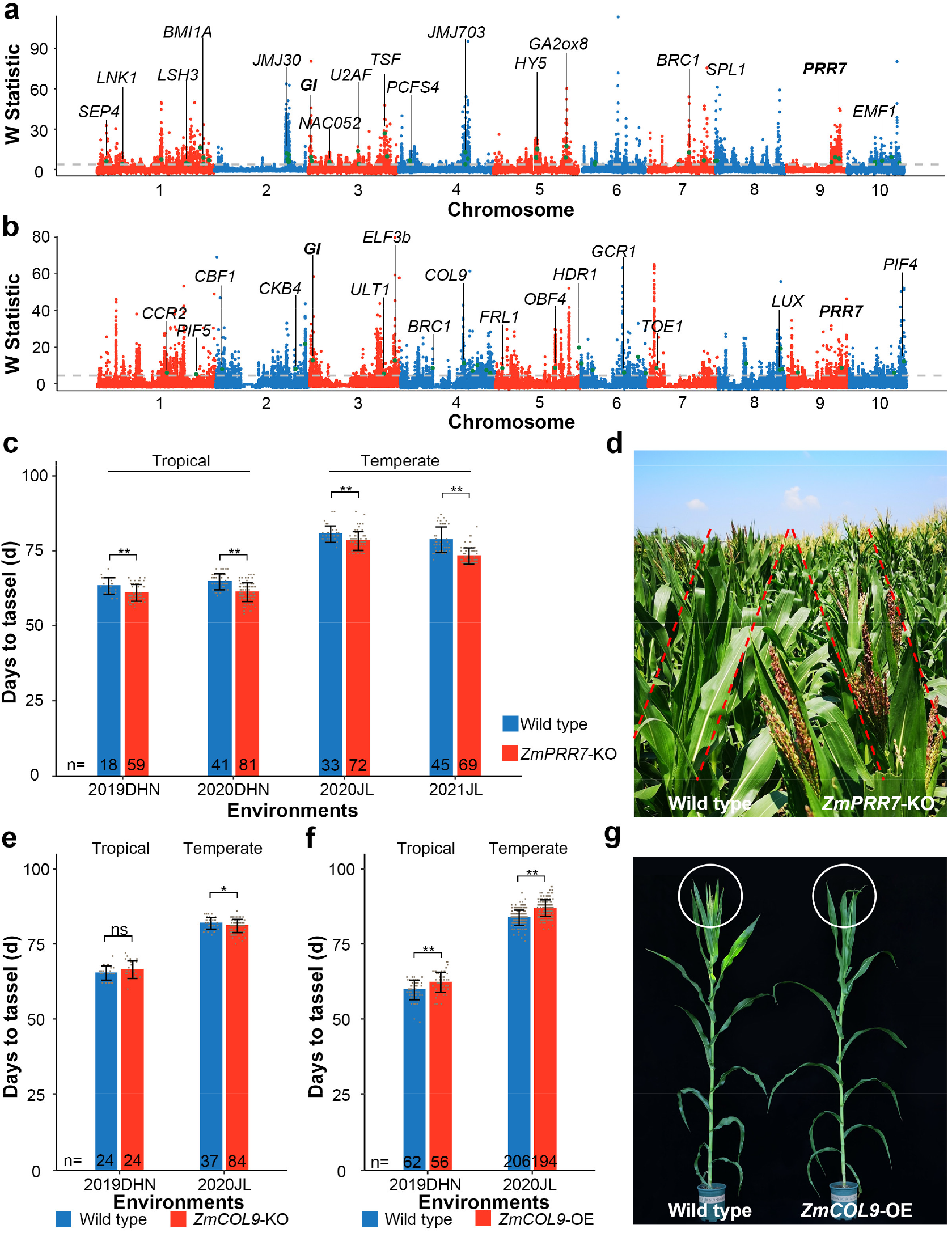
Local adaptation in teosinte and maize. Genome-wide selection signals (W statistic reflecting smoothed XP-CLR score) between **a**, *mexicana* and *parviglumis*; and **b**, temperate and tropical maize. The horizontal grey dashed line represents the top 5% cutoff. Genes associated with flowering time and floral development in maize, rice and *A. thaliana* are marked with green points. **c**, Days to tassel of wild type and *ZmPRR7* knock out (KO) mutants under tropical (Hainan province 2019 and 2020; China; E109°, N18°) and temperate (Jilin province 2020 and 2021; China; E125°, N44°) environments. **d**, *ZmPRR7* KO mutants showed earlier flowering relative to wild type. The picture was taken in Jilin province 2020 at 77 d after planting. **e**. Days to tassel of wild type and *ZmCOL9* KO mutants under tropical and temperate environments. **f.** Days to tassel of wild type and *ZmCOL9* over-expression (OE) mutants under tropical and temperate environments. **g**, *ZmCOL9* OE mutants showed later flowering relative to wild type. The picture was taken in Jilin province in 2020 at 78 d after planting. ns indicates non-significant difference between mutants and wild type by two-sided t-test at *P*-value = 0.05, * indicates *P*-value < 0.05, ** indicates *P*-value < 0.01.

Selection for variants that promote early flowering enabled maize to break day-length restrictions and facilitated the spread of maize across a broad geographical range^82^, and the alleles involved in flowering time are also a major target of highland landrace adaptation^83^. Experimental data in maize^84^ and from orthologs^85^ in other species shows that at least 51 genes associated with highland and 61 genes associated with high latitude adaptation were involved in flowering time pathways (Supplementary Fig. 20 and Supplementary Table 22), respectively. For example, the genes *GI* and *PRR7*, both known to participate in the circadian clock pathway in *Arabidopsis* and rice^86–89^, show evidence of selection both in *mexicana* and temperate maize. Tracking the flowering time pathway, we found temperate maize has more genes under selection in the photoperiod pathway (eight in temperate maize, five in *mexicana*; Supplementary Table 22), which may be a signal of adaptation to changing latitude.

To validate the utility of the selection scan approach, we tested the function of *ZmPRR7* (*Zm00001d047761*), which shows convergent patterns in maize and teosinte, and the maize-specific candidate *ZmCOL9* (*Zm00001d051684*) that is involved in the photoperiod pathway. Mutants of these two genes were obtained from a CRISPR/Cas9-based high-throughput targeted mutagenesis library^90^. The mutant allele of *ZmPRR7* is a 5.8-Kb deletion in the gene region that leads to the total loss of protein function. Plants harboring the mutant allele exhibit significantly earlier flowering than the wildtype in both tropical and temperate environments (Fig. 3c, d and Supplementary Fig. 21). The loss-of-function allele of *ZmCOL9* includes a 5 bp deletion/1bp insertion in the intron and a 2 bp deletion/4 bp deletion in the 3^rd^ exon (Supplementary Fig. 22a, d) that result in premature translation termination. In a tropical environment (Hainan; China; E109°, N18°), *ZmCOL9* knockout mutants showed no difference in flowering time compared to the wild type (Fig. 3e and Supplementary Fig. 23b, e) but overexpression plants exhibit a later flowering phenotype (Fig. 3f and Supplementary Fig. 23a, b). In contrast, when planted in a temperate environment (Jilin; China; E125°, N44°), the *ZmCOL9* knockout mutants flowered earlier (Fig. 3e and Supplementary Fig. 22c, f) and the overexpression lines flowered later than the wild type (Fig. 3f, g and Supplementary Fig. 23c, d). These results confirmed the key roles for both *ZmPRR7* and *ZmCOL9* in regulating flowering time and contributing to the adaptation of highland teosinte and modern maize.

## Discussion

The twin projections of increasing human population and decreasing suitable farmland highlight the challenge breeders face in producing high crop yields, and this has motivated an increasing interest in crop wild relatives as sources of genetic diversity for improvment^91,92^. Here, we present a high-resolution genetic variation map that greatly expands the publicly available genetic sequence information for the genus *Zea*.

We provide the first complete picture of the phylogeny and demography of the genus *Zea* using genome-wide data, including both divergence times and effective population sizes of *Zea* species. We reaffirm several aspects of the phylogeny of *Zea*, but our data identify a number of new features, including the likely subspecies status of *nicaraguensis*, the short divergence times between the perennial taxa, and the relatively young age of the genus. We caution that our divergence estimate for *Tripsacum* may be underestimated because of the difficulty of mapping short reads from divergent genomes, however, and that high-quality *Tripsacum* and teosinte reference genomes will be essential to better answer this question^93^.

Our broad sampling of the genus allows us to take advantage of population genetic tools to identify candidate genes involved in adaptation across both long and short time scales. We find evidence of convergent adaptation of highland teosinte and high-latitude maize, exemplifying the utility of studying variation in wild relatives to identify genes important in crops. Finally, we validate these approaches by using genome editing to knock out two candidate flowering time genes. All data and results of this work have been integrated into the ZEAMAP database^94^ for easy query and retrieval.

It is particularly noteworthy that our work identifies a vast trove of genetic variation absent in cultivated maize and even in its closest wild relative *parviglumis*. Our functional analysis of candidate adaptation genes clarifies the great potential in the utilization of the wild relatives of maize in identifying novel alleles or highlighting potential genes for subsequent editing, potentially accelerating modern genetic improvements^95^. The data and discoveries presented in this study provide the foundation for the use of crop wild relative resources for breeding in the face of increasing human populations and decreasing farmland.

## METHODS

### Samples and whole genome resequencing

A total of 237 teosinte accessions from CIMMYT, USDA and collaborators were obtained, consisting of 90 *mexicana*, 79 *parviglumis*, 20 *diploperennis*, 15 *perennis*, 15 *luxurians*, 13 *nicaraguensis*, five *huehuetenangensis* according to morphological classification (Supplementary Table 1, 2). Two *Tripsacum dactyloides* were obtained from Dr. Fajia Chen’s lab (Henan Agricultural University, China). Young leaves from one individual of each accession were used for DNA extraction for sequencing using the Illumina HiSeq3000 platform (150-bp paired-end), conducted by BGI (Shenzhen, China) and NovaSeq6000 platform (150-bp paired-end), conducted by Novogene (Sacramento, USA). DNA sequencing data of 507 cultivated maize were downloaded from the NCBI SRA database (PRJNA531553; Supplementary Table 1).

### Read mapping and SNP calling

Raw reads of teosinte were first processed using FastQC (v0.11.3; http://www.bioinformatics.babraham.ac.uk/projects/fastqc/). Trimmomatic^96^ (v0.33; HiSeq3000 platform; LEADING:3 TRAILING:3 SLIDINGWINDOW:4:15 MINLEN:36) and fastp^97^ (v0.19.4; NovaSeq6000 platform; -g -l 36) were used to remove poor-quality base calls and adaptors. Reads of teosinte and maize were then aligned to the B73 reference genome^98^ (v4) using Bowtie2^99^ (v2.1.0; --very-fast). Unique mapped reads were sorted and indexed using Picard (v1.119; http://broadinstitute.github.io/picard/). SAMtools^100^ (v1.3.1) and UnifiedGenotyper from GATK (v3.5; https://software.broadinstitute.org/gatk/) were used to estimate the variant calling file for each individual. Hard filtering of the individual SNP calls was carried out with mapping quality (MQ ≤ 20.0), and thresholds set by sequencing coverage based on minimum coverage (DP ≤ 5) and maximum coverage (DP ≥ 200). Then, variants from the 237 teosinte and 507 maize were combined by GATK CombineVariants to a single variant calling file. To confirm if unknown variants were discarded reference genotypes in individual calls, we recalled these sites and replaced them with reference genotypes if they had supported reads. Finally, sites with a missing rate higher than 75% in all samples were excluded. To validate the accuracy of SNPs called from resequencing data, 224 sites in 80 accessions were selected for Sanger sequencing (Supplementary Table 3).

### Population structure classification, principal component analysis and phylogenetic tree construction

We evaluated patterns of population structure using a set of SNPs filtered to remove multi-allelic loci and SNPs with a minor allele frequency < 0.05 (--maf 0.05 -biallelic-only) using PLINK^101^ (v1.9). We then ran admixture^102^ for different values of the number of clusters (K) from 2 to 20 (--cv = 10; v1.3.0). Each individual with admixture components < 0.6 was classified as ‘Teosinte (mix)’ or ‘maize (mix)’. We performed PCA analysis using this same set of SNPs with GCTA^103^ (v1.26) recording the first 10 components (--pca 10). We annotated SNPs with a missing data rate less than 0.7 in teosinte and maize with SnpEff (v4.1g; http://snpeff.sourceforge.net/index.html) using the first transcript of B73 v4 genes. We then used synonymous and noncoding SNPs to construct a simple phylogenetic tree with SNPhylo^54^ (v20140701) using default parameters and visualized the tree with iTOL^104^.

### Species tree analysis

Species delimitation and species trees were inferred using BPP^53^ (model A11; v4.1.4). We used the following samples in BPP: three tropical maize, three *parviglumis*, three *mexicana*, three *nicaraguensis*, three *diploperennis*, three *perennis*, three *luxurians*, two *huehuetenangensis* and two *Tripsacum dactyloides* (Supplementary Table 1). Low-quality base calls and adaptors from raw reads of *Tripsacum dactyloides* were removed using Trimmomatic, and the remaining sequences were aligned to the B73 v4 reference genome with Bowtie2 as described above. The consensus base was estimated from the uniquely mapped reads using ANGSD^105^ (v0.930). Using the B73 annotation, we randomly selected 2,000 coding sequence genes to estimate the species delimitation and species tree. The prior distribution of ancestral population size (θ) and divergence time from the root (τ) followed an inverse-gamma (IG) prior with means of 0.005 IG (3 0.01)) and 0.75 (IG (3 1.5)), respectively. The consensus of A11 species trees was visualized using DensiTree^106^ (v2.2.6).

### Imputation and demographic estimation

SNPs in the 237 teosinte and 507 maize were imputed with BEAGLE^107^ (v4.0), respectively. Divergence times within teosinte and the effective population size of each teosinte were estimated using BPP (A00 model) and MSMC2^57^ (v2.1.1). The topological tree in BPP (A00 model) was fixed as the species tree with highest posterior probability (A11 model) estimated from the above species tree analysis. Sequences used in the A11 model were applied to estimate the effective population size and divergence time using priors as above. In MSMC2, four haplotype models were applied (Supplementary Table 1). The mutation rate used in BPP (A00 model) and MSMC2 was 3E-08^108^.

### ABBA-BABA and divergence-based introgression polarization test

We used Patterson’s D statistic^109,110^ to test for introgression between teosinte. Assuming *Tripsacum dactyloides* as the outgroup (O), we assessed D statistics for the tree (((P1, P2), P3), O), P1/P2/P3 representing different taxa in *Zea* (autotetraploid *perennis* was excluded). The number of ABBA and BABA in each block were calculated in ANGSD (-blockSize 10000). To overcome the problem of non-independence within the sequence, a block-jackknifing procedure was used to test for statistical significance. To estimate the directions of introgression, consensus base was estimated from the uniquely mapped reads using ANGSD for representing individuals in different taxa of *Zea* and *Tripsacum* (eight taxa in total). The whole-genome consensus files from different taxa were then concatenated into multiple sequence alignment files by different chromosomes. Finally, this eight-taxon alignment was pruned to contain four taxa according to each test as suggested in Supplementary Fig. 6 and divided into a 5,000bp windows, which were used as the input of DIP^111^.

### Linkage disequilibrium, nucleotide diversity and *F*_ST_ calculation

Linkage disequilibrium (*r*^2^) of *nicaraguensis* (14), *luxurians* (14), *diploperennis* (20), *perennis* (15), *huehuetenangensis* (five), *mexicana* (81), *parviglumis* (70), and maize (507) were estimated for all bi-allelelic SNPs within 500Kb (--geno 0.5 --maf 0.05 --biallelic-only --snps-only) using PLINK. Nucleotide diversity of *nicaraguensis* (14), *luxurians* (14), *diploperennis* (20), *perennis* (15), *huehuetenangensis* (five), *mexicana* (81), *parviglumis* (70) and maize (randomly selected 110 individuals) was calculated using ANGSD (v0.930, -doMaf 1 -doMajorMinor 1 -uniqueOnly 1 -minMapQ 30 -minQ 20 -GL 2 -fold 1 -win 5000 -step 5000). Differentiation (*F*_ST_) between maize and teosinte with five randomly selected samples was estimated in VCFTools^112^ (v0.1.16; --fst-window-size 5000).

### Taxon-specific SNPs, InDels and k-mer analysis

SNPs and InDels found only in one specific taxon in *Zea* in at least two individuals were regarded as taxon-specific SNPs. The longest transcripts of each gene in the B73 annotation and a recent atlas of cis-regulatory elements^59^ were used to annotate variants. K-mers of teosinte and maize were counted using Jellyfish^113^ (v2.3.0; -m 31). K-mers unique to each taxon that appeared at least two times were obtained with sourmash^114^ (v3.2.0; --scaled 1000).

### Transposon element analysis

RepeatExplore2^115^ was used to identify repeat clusters of each taxa of *Zea* (two samples were randomly selected from each taxon). Clusters were further annotated by applying RepeatMasker (http://www.repeatmasker.org/; v4.1.0; -species maize). Reads were mapped to the above repeat clusters by using BWA-MEM^116^ (v0.7.10), and the number of mapped reads in each repeat clusters were calculated with SAMTools. Abundance of the repeat elements between samples were normalized by their sequenced library size.

### Inversion calling

Localized heterogeneity across chromosomes was identified using lostruct^68^ in windows containing 10,000 SNPs. The most related 5% of windows in each chromosome around one of the four outliers (maximum, minimum MDS1 or MDS2) were regarded as candidate inversions and were genotyped using invClust^117^ (v1.0) with B73 as the reference state. Genotypes of the candidates were confirmed via PCA of the SNPs in the corresponding region. Only taxa with three clearly different haplotypes identified by PCA were regarded as true inversions. Candidates near the centromeres were filtered out. Centromere information was obtained by combining locations from entire in the NAM population^63^.

### Genome-wide association analysis

SNPs from *mexicana* were obtained from the imputed teosinte panel according to the name of samples, and then population structure was calculated with admixture (v1.3.0; --cv=10; K=1, 2, 3, 4, 5). The K value with the lowest CV (K=2) was used in downstream analysis. Estimation of the kinship matrix and association analyses using the compressed MLM were performed using TASSEL3^118^ (v3.0.174), with a *P*-value cut off set to 1/N (N = the number of tested SNPs). Latitude and longitude information was obtained from Dr.Suketoshi Taba’s lab. Global soil properties used as phenotypes for the GWAS were extracted using the R package ncdf4 (v1.16; http://cirrus.ucsd.edu/~pierce/ncdf/) from the Global Soil Dataset for Earth System Modeling^69^, a comprehensive database with eight layers to the depth of 2.3m (0-0.045, 0.045-0.091, 0.091-0.166, 0.166-0.289, 0.289-0.493, 0.493-0.829, 0.829-1.383 and 1.383-2.296 m). Soil properties were clustered using the R package clValid^119^, which tested hierarchical, k-means and k-medoide in combination with 2-40 clusters to find the best method and cluster numbers. GWAS were performed on a subset of nine features identified by hierarchical cluster analysis (Supplementary Fig. 16).

### Identification of adaptive regions in non *Zea mays* taxa

Whole genome adaptive genetic variation between different non *Zea mays* taxa and maize were estimated by calculating their *F*_ST_ value in VCFTools (--fst-window-size 5000). Under each comparison, all available teosinte and maize samples were used. We then Z-transformed the *F*_ST_ in each window, windows with Z*F*_ST_ values exceeding the 95^th^ percentile of the whole genome were declared as candidate adaptive regions. GO enrichment analysis was conducted using PANTHER with default parameters^120,121^ and visualized with GlueGo^122^.

### Selective sweeps in teosinte and maize

Whole genome scanning for regions of teosinte elevation adaptation and maize temperate adaptation was implemented by a mixed method. First, two genetic maps were obtained from a B73 x Teosinte population^123^ and a maize B73 x By804 population^124^, and the physical locations were converted to coordinates of the B73 v4 reference sequence using CrossMap^125^ (v0.2.9). The genetic distance between SNPs in *mexicana* and *parviglumis* were then calculated based on the B73 x Teosinte genetic map, while the distance in temperate maize and tropical maize were calculated based on the B73 x By804 genetic map. Genetic distances between SNPs located between the genetic markers were assigned based on their physical distance. The likelihood of multi-locus allele frequency differentiation between two tested populations was modeled using XP-CLR^126^ (v1.0; -w1 0.005 100 1000 -p0 0.7) in both the teosinte group (*mexicana*, with *parviglumis* as the reference) and the maize group (temperate maize, with tropical maize as the reference). Finally, we applied a spline-window method (GenWin^127^ v0.1; smoothness = 100) to smooth the results. The top 5% of genomic region with the highest W statistic in *parviglumis* and *mexicana* were regarded as candidate teosinte altitude adaptation regions and the top 5% of the W statistic regions in temperate and tropical maize were regarded as candidate maize temperate adaptation regions. Enrichment analysis between candidate teosinte altitude adaptation regions and maize temperate adaptation was conducted using the shuffle function (-excl -noOverlapping) in BEDTools^128^ (v2.25.0). Genes, including the promoter and 2kb upstream, that overlapped with the regions identified above were regarded as candidate adaptive genes.

### RNA-seq sampling, library construction and data analysis

The base tissues of V5 stage shoots (1-2 cm) of maize (five tropical maize; five temperate maize) and teosinte (three *parviglumis*; three *mexicana*) were sampled for mRNA and total RNA extraction. Both mRNA and total RNA samples were used for library preparation according to Illumina strand-specific library construction protocols. Paired-end libraries were sequenced using a mixture of platforms (Hi-Seq3000, x10, NovaSeq) with 150 cycles. Raw reads were filtered to remove the poor-quality base calls and adaptors specifically for each platform (NovaSeq: fastp -g -l 36; x10: fastp -l 36; Hi-Seq3000: Trimmomatic LEADING:3 TRAILING:3 SLIDINGWINDOW:4:15 MINLEN:36). Reads were then aligned to the B73 reference genome (V4) using TopHat2^129^ (v2.2.1) and read counts for each gene were calculated using htseq-count^130^ (v0.9.1). Finally, differentially expressed genes were identified between tropical and temperate maize, as well as between *parviglumis* and *mexicana*, using DESeq2^131^ (v1.10.1) with absolute fold change higher than 1 and *P*-value < 0.05.

### Functional validation of *ZmPRR7* and *ZmCOL9*

Mutants of *ZmPRR7* and *ZmCOL9* were generated from a high-throughput genome-editing design^90^. In brief, line-specific sgRNAs were filtered based on the assembled pseudo-genome of the receptor KN5585, and a double sgRNAs pool (DSP) approach was used to construct vectors. The vectors were transformed into the receptor KN5585, and the targets of each T0 individual were assigned by barcode-based sequencing. The genotype of gene-editing lines was identified by PCR amplification and Sanger sequencing using target-specific primers (Supplementary Table 23).

Transgenic lines generated with DNA fragments of *ZmCOL9* driven by the *ZmUbi* promoter were created using the modified binary vector pCAMBIA3300. Immature zygotic embryos of maize hybrid HiII (B73 x A188) were infected with *A. tumefaciens* strain EHA105 harboring the binary vector based on the published method for *ZmCOL9*^132^. Transgenic plants were identified by qRT-PCR as well as tests for herbicide resistance and the presence of the bar gene. Flowering-time phenotypes of mutants and transgenic plants of *ZmPRR7* and *ZmCOL9* were investigated in Jilin province (E125°, N44°) and Hainan province (E109°, N18°).

## Data availability

DNA- and RNA-sequencing reads from this study were deposited in the NCBI Sequence Read Archive with the accession number of PRJNA641489, PRJNA816255, PRJNA816273 and PRJNA645739, respectively. The SNP data can be downloaded from https://ftp.cngb.org/pub/CNSA/data3/CNP0001565/zeamap/02_Variants/PAN_Zea_Variants/Zea-vardb/.

## Code availability

All custom scripts used in this study are available at https://github.com/conniecl/Zea_genus.

## ACKNOWLEDGMENTS

We would like to thank Dr. Suketoshi Taba from CIMMYT for providing teosinte materials. We would like to thank Dr. Jiafa Chen from Henan Agricultural University for providing *Tripsacum dactyloides*. We would like to thank Mr. Hao Liu from the high-throughput computing platform of National Key Laboratory of Crop Genetic Improvement. We would like to thank Andi Kur from North Carolina State University for drawing the picture of teosinte morphological characteristics. We also thank Taylor AuBuchon-Elder and Sally Fabbri for growing and supplying germplasm. This research was supported by the National Key Research and Development Program of China (2020YFE0202300) and National Natural Science Foundation of China (U1901201, 31771879), as well as the US National Science Foundation (grants 1546719,1822330) and USDA Hatch project CADPLS2066H.

## AUTHOR CONTRIBUTIONS

J.Y., J.R.-I. and N.Y. designed and supervised this study. Y.P., W.L., A.P., B.C., J.B., R. R.-A., R.S., J.Y., Q.Z., S.W., S.G., Y.W., Y.L., C.J., M.D., M.J., J.L., L.J., Y.Y., M.Z. and X.Y. prepared the materials. X.Z. provided the variant calling pipeline. J.L. performed the Sanger validation of SNPs. W.W uploaded the SNPs and InDels to the database. L.C. and J.L. analyzed the data. M.J., X.L., L.Q., Y.Y., and X.Y performed genetic transformation and mutant validation. L.C., M.J., N.Y., M.H., A.R.F., M.L.W., J.R.-I. and J.Y. prepared the manuscript.

## COMPETING FINANCIAL INTERESTS

The authors declare no competing financial interests.

## Notes

### Competing Interest Statement

The authors have declared no competing interest.

